# Short Linear Motifs as a General Organizing Principle of the Nuclear-Receptor Proximal Interactome

**DOI:** 10.64898/2026.08.06.743235

**Authors:** Jimmy K. Eng, Lilliana Radoshevich, Michael E. Wright

**Affiliations:** University of Washington Proteomics Resource, 850 Republican Street, Seattle, WA 98109, USA; Department of Immunology and Genomic Medicine, National Jewish Health, 1400 Jackson Street, Denver, CO 80206, USA; Department of Molecular Physiology and Biophysics, Carver College of Medicine, University of Iowa, 51 Newton Road, Iowa City, IA 52242, USA

## Abstract

The AR-interactome comprises ∼1,000 androgen receptor-interacting proteins (AR-IPs), yet how a single receptor engages so many partners across compartments remains mechanistically unclear. We integrate proximity-labeling quantitative mass spectrometry across cytosolic, microsomal, and nuclear compartments of LNCaP prostate cancer cells and resolve 4,751 AR-proximal interacting proteins (AR-PIPs), more than four times the size of the AR-interactome. Anchoring on the LXXLL coactivator recognition motif, LXXLL motifs are systematically depleted in AR-PIPs after length control, consistent with low-affinity, transient engagement at the AR AF-2 charge clamp. LXXLL-bearing AR-PIPs include AR itself and canonical AR coactivators altered by amplification, deletion, or motif-spanning mutations in metastatic and castration-resistant prostate cancers. AR-V7, which lacks AF-2, retains LXXLL-depleted Mode 1 partners and loses the LXXLL-enriched Mode 2 cloud, thereby validating a two-mode engagement framework for nuclear receptor-proximal interactomes.

## Introduction

The androgen receptor (AR) is a ligand-activated transcription factor that orchestrates male development, reproductive physiology, and prostate cancer progression, with broader roles in non-reproductive tissues [1,2]. Four decades of biochemical and proteomic efforts have cataloged nearly 1,000 AR-interacting proteins (AR-IPs), collectively known as the AR-interactome, spanning coactivators, corepressors, chromatin remodelers, pioneer factors, and post-translational enzymes [1,3]. Despite this catalog, two questions remain unresolved. How does AR physically engage the much larger spatial neighborhood that proximity labeling consistently captures beyond the curated AR-interactome [4,5,6], a pattern also seen for other steroid receptors including ER [7] and GR [8]? And what structural features of the AR ligand-binding domain (LBD) allow a single receptor to sample so many partners?

The companion extranuclear [9] and nuclear [10] AR-proximal interaction network (AR-PIN) atlases established that the proximal AR-interactome, aka AR-proximal interacting proteins (AR-PIPs), is a temporally resolved continuum across the cytosolic, microsomal, and nuclear compartments, with any single compartment exceeding the AR-interactome [3]. Here, we integrate the three compartments and ask which motif architecture organizes the AR-proximal proteome. Short linear motifs (SLiMs) are brief amphipathic stretches of three to ten residues that mediate low-affinity, competitive, and transient interactions in disordered protein regions [11,12]. The LXXLL nuclear receptor box, first described as the coactivator recognition motif in p160 family members [13,14], and the AR-specific FXXLF (FQNLF) motif [15], engage a single hydrophobic groove on the AR LBD, the activation function 2 (AF-2) surface, framed by a charge clamp at Lys720 and Glu897 [16,17,18].

Full-length cryo-EM [19] shows that agonist-bound AR forms an intramolecular FQNLF-to-AF-2 lock within each monomer, and the AR dimer displays this lock in both subunits. At the same time, SRC-3 binds the AR N-terminal domain (NTD) through structured bHLH/PAS, S/T, and HAT contacts independent of LXXLL. Bona fide AR coregulators therefore engage the NTD through high-affinity structured-domain contacts [20], while the AF-2 charge clamp transiently and competitively samples the proteome-wide LXXLL pool (Mode 2) when intramolecular FQNLF has not yet locked it.

Here, we test this two-mode framework directly. Using MotifHunter, a sequence-scan algorithm developed for this study, we split the canonical AR-PIP sets from each compartment by LXXLL motif content and controlled for protein length. We anchor the framework to clinical disease by integrating cBioPortal and COSMIC mutation data on LXXLL-bearing AR-PIPs in human prostate cancers. Lastly, we exploited the AR-V7 splice variant, which lacks AF-2 [4], as a natural ΔAF-2 experiment, predicting the loss of the Mode 2 LXXLL-enriched cohort.

## Results

### Size of the AR-PIP universe across compartments

To define the AR-PIP universe, we used PL-qMS [21] to resolve the proximal AR-interactome across cytosolic, microsomal, and nuclear compartments in LNCaP prostate tumor cells at 6 time points in response to androgens [9,10] (Figure 1A). Mapping the 989-member AR-interactome (BioGRID, NCBI, and published interaction databases; Methods) against these datasets showed that 76.5% (757 of 989) of AR-IPs were recovered as AR-PIPs, and that PL-qMS detected 87.6% (866 of 989) across at least one compartment (Supplementary Fig. 1). Per-compartment recovery was highest in the nucleus (664 members, 67.1%) relative to microsomal (634, 64.1%) and cytosolic (564, 57%) compartments (Figure 1B), consistent with the chromatin-directed Launonen et al (2021) ChIP-SICAP chromatome benchmark [9,10]. Because proximity labeling reports the full molecular neighborhood rather than a curated set of functional partners, this AR-PIP universe comprises both direct functional interactors and proximal bystanders; the motif grammar we read across it is therefore a property of the entire proximal population, not of a pre-selected interactor list.

**Figure 1.**
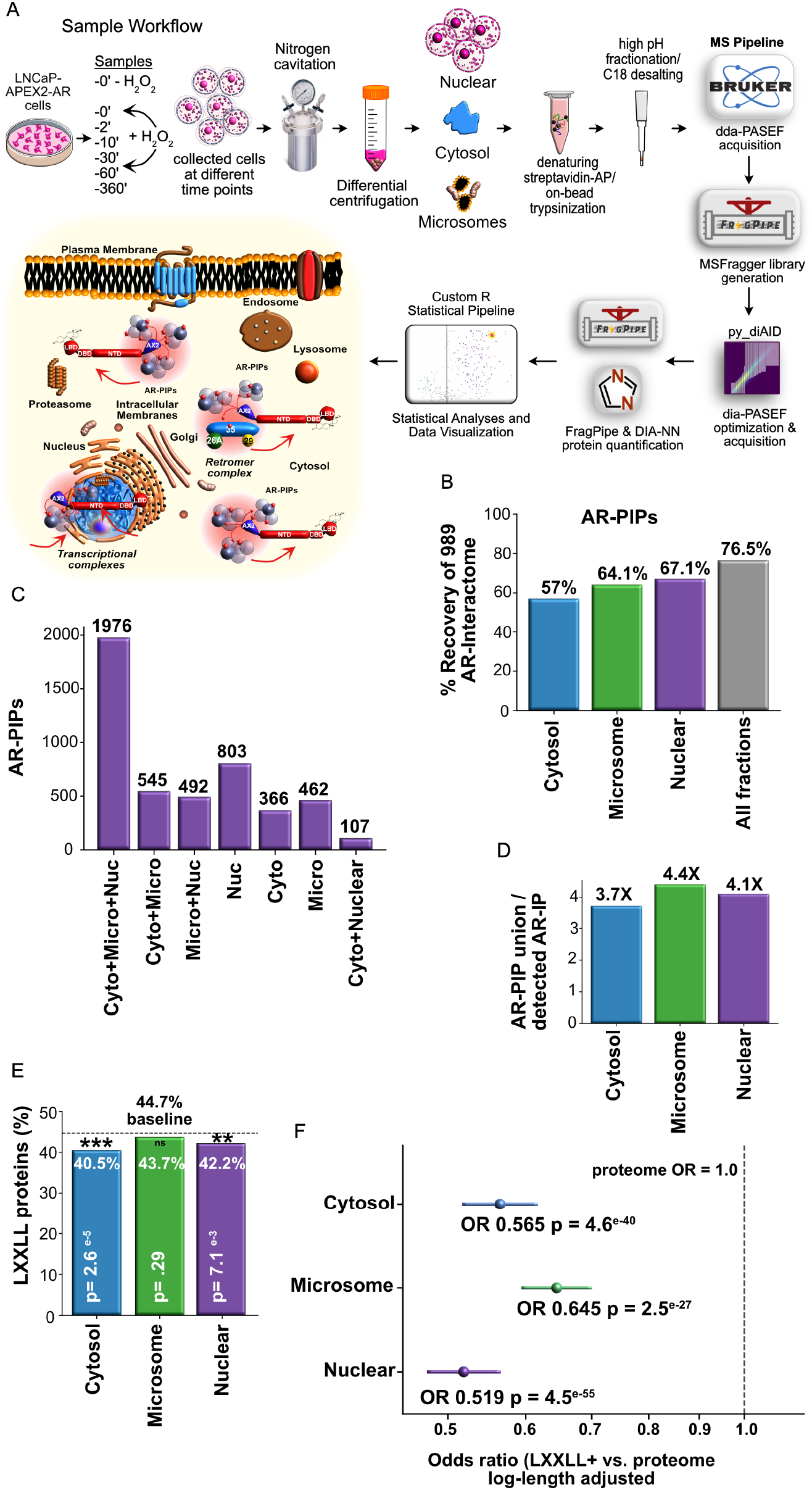
Cross-compartment AR-PIP universe expands the AR-interactome four to five fold. (A) Experimental workflow for compartment-resolved APEX2-AR proximity labeling in LNCaP prostate cancer cells across six androgen time points (0, 2, 10, 30, 60, 360 min). Cells are lysed by nitrogen cavitation, fractionated by differential centrifugation into cytosolic, microsomal, and nuclear compartments, and streptavidin-purified biotinylated proteins are analyzed by dda-PASEF plus dia-PASEF mass spectrometry. (B) Percent recovery of the 989-member curated AR-interactome (AR-IP catalog) as AR-PIPs per compartment and pooled across all fractions (denominator 989). (C) AR-PIP universe by set membership across the three compartments (n = 4,751 total unique AR-PIPs, three-way intersection n = 1,976). (D) Per-compartment expansion factor (AR-PIP union over detected AR-IP catalog members). (E) LXXLL-positive prevalence in AR-PIPs per compartment, relative to the whole-proteome baseline of 44.7 percent (UniProt 2023_10, n = 20,211 reviewed entries). Significance from chi-square test. n.s. not significant, ** p < 0.01, *** p < 0.001. (F) Length-controlled logistic regression odds ratios for LXXLL-positive status by compartment membership against the whole-proteome baseline (adjusted for log protein length). Reference OR = 1 dashed line. All p < 1e-26. See also Supplementary Fig. 1 and Figure 2. Source data, Source Data (sheets Fig_1B through Fig_1F); panel A is a schematic.

The AR-PIP universe extends well beyond the AR-interactome, encompassing 2,994, 3,475, and 3,378 AR-PIPs in the cytosolic, microsomal, and nuclear compartments, respectively (Supplementary Data 1, sheet ARPIP_Universe). AR-PIPs union across the three compartments resolved 4,751 unique proteins, with 1,976 shared across all three compartments (Figure 1C). Compartment-specific AR-PIPs included 366, 462, and 803 cytosolic, 462 microsomal, and 803 nuclear proteins, respectively. The remaining shared AR-PIPs were distributed across pairwise intersections (Figure 1C). The overlap of AR-PIPs across the cellular cartography indicates that a stable core of proximal AR interactors is present in ligand-activated AR. In contrast, compartment-specific AR-PIPs contribute to the spatial context that is unresolvable by a single-compartment proteomic experiment.

The AR-PIP universe is more expansive than the AR-interactome, with the AR-PIP union exceeding the detected AR-interactome by approximately four-fold in each compartment (*i.e.*, cytosolic 3.7-fold, microsomal 4.4-fold, nuclear 4.1-fold) (Figure 1D). Overall, the AR-PIP universe expands the AR-interactome catalog by approximately five-fold, and this expansion is not an artifact of PL-qMS sensitivity. The AR-interactome was assembled from diverse biochemical and proteomic technologies, including yeast two-hybrid, affinity purification, RIME, and earlier proximity-labeling studies [3,5,6,22,23]. The scale of the expansion raises a structural question. What features of AR, a single receptor molecule, enable it to engage so many partners?

### Length-controlled LXXLL depletion across the AR-PIP universe

The LXXLL motif is the canonical nuclear receptor coactivator recognition element [13,14], a short amphipathic helix engaged by the AR AF-2 charge clamp alongside the related FXXLF and WXXLL variants [15,17]. Because LXXLL motifs are distributed across thousands of proteins in the human proteome, they are a plausible structural rationale for the AR-PIP expansion. If LXXLL engagement at the AR AF-2 charge clamp drives AR’s broad proximal proteome, LXXLL-bearing proteins should be over-represented among AR-PIPs relative to the proteome baseline.

Testing the LXXLL hypothesis at the proteome scale required a tool that could scan large protein databases for multiple short linear motifs in a single pass and integrate the resulting motif assignments directly into our compartment-resolved AR-PIP analysis pipeline. Pattern matching against UniProt [24] has been performed by prior tools, including EPSLiM for nuclear receptor short linear motifs [12]. We built MotifHunter as a lightweight, open-source workflow tool that supports concurrent regex-based scans for LXXLL, FXXLF, and WXXLL nuclear receptor box motifs across whole proteomes, with output that interfaces directly with HGNC-canonical AR-PIP aggregation and downstream statistical analysis (see the Key Resources supplementary table; Methods). The extended motif and domain scan built on this workflow, and the Predictive Proximal Proteome Database assembled from it, are presented in the companion resource [25].

To test this prediction, we applied MotifHunter to scan canonical Swiss-Prot UniProt sequences (UniProt 2023_10; 20,211 reviewed entries) for the LXXLL, FXXLF, and WXXLL motifs and split the per-compartment AR-PIP sets by motif content. Across the human proteome, 44.7% of canonical sequences contain at least one LXXLL motif. Within the AR-PIP populations, the raw LXXLL prevalence was 40.5% in cytosolic, 43.7% in microsomal, and 42.2% in nuclear AR-PIPs (Figure 1E). The cytosolic and nuclear AR-PIP sets are significantly depleted of LXXLL motifs relative to the proteome (chi-square p = 2.6 × 10⁻⁵ and 7.1 × 10⁻³, respectively), and the microsomal set is statistically indistinguishable from baseline (p = 0.29). The depletion of the LXXLL motif among AR-PIPs was a surprise finding and runs opposite to the intuitive expectation that AR’s coactivator recognition mechanism should be reflected in LXXLL enrichment among AR-PIPs.

To test whether the depletion reflects compartment biology rather than confounding due to protein length, we compared the protein-length distribution of each AR-PIP set against the proteome baseline. The expected frequency of LXXLL motifs in a sequence scales approximately as 5L/20⁵, where L is the amino acid length, so length differences between compartments and the proteome baseline could mechanically inflate or deflate the prevalence of LXXLL motifs. Unexpectedly, the AR-PIP sets are systematically biased toward larger proteins (*i.e.*, median 561 to 633 amino acids versus 417 in the proteome; fraction below 150 amino acids approximately 2% versus 11%; Figure 2A). The small protein confound, which would have predicted mechanical depletion of LXXLL, runs in the opposite direction of the observed depletion. Binning LXXLL-positive prevalence directly by protein length confirms that within every matched length window, the cytosolic, microsomal, and nuclear AR-PIPs each carry a lower LXXLL-positive fraction than the whole proteome (Figure 2B), establishing the depletion as a within-bin property rather than a modeling artifact.

**Figure 2.**
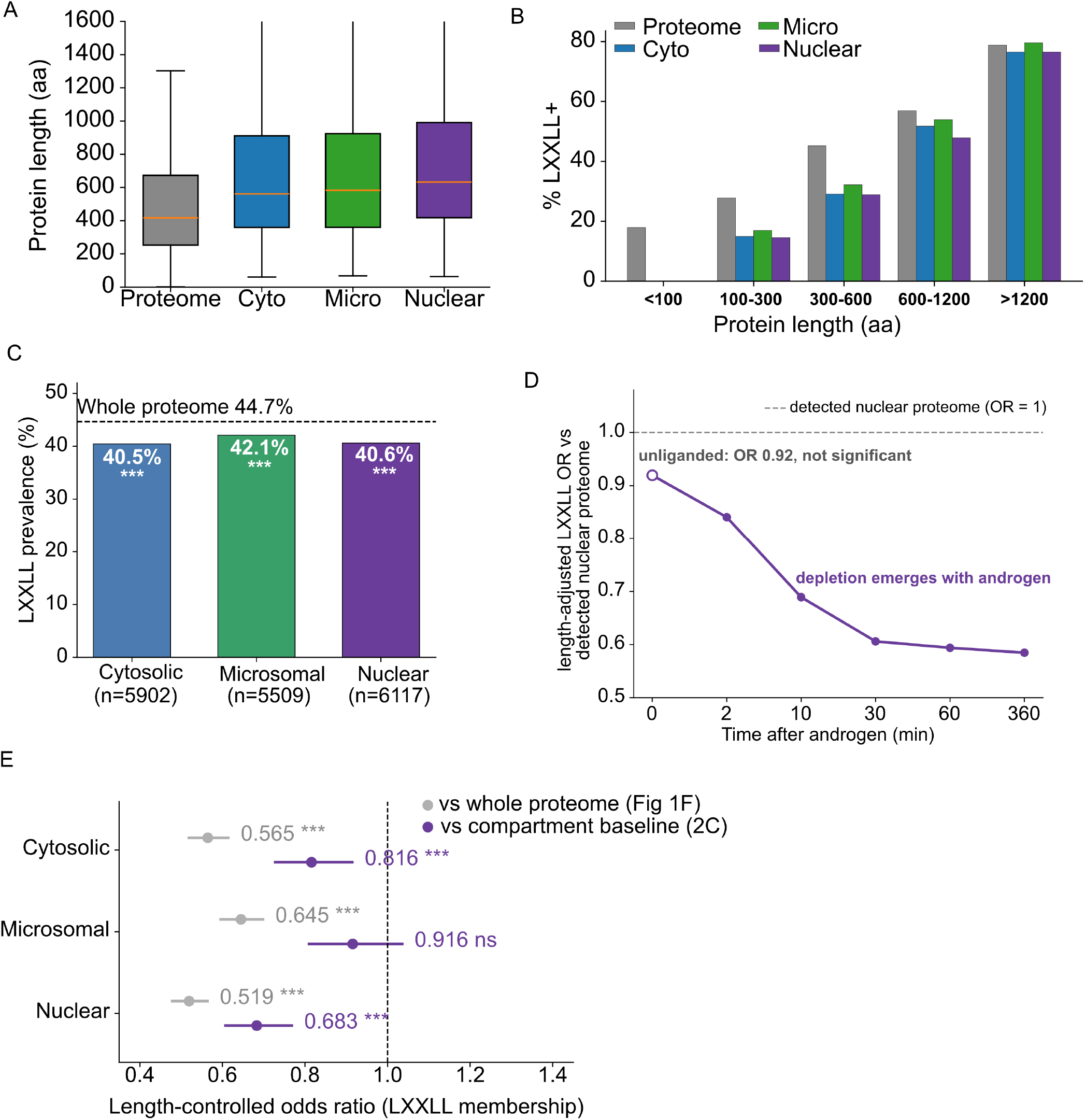
Protein length, LXXLL prevalence by length, and compartmental LXXLL baseline controls. (A) Cumulative protein length distribution for cytosolic, microsomal, and nuclear AR-PIPs compared with the entire UniProt 2023_10 proteome (n = 20,211 reviewed entries). Median lengths, cytosolic 561, microsomal 583, nuclear 633 amino acids; proteome 417 amino acids. The fraction of proteins under 150 amino acids is approximately 2 percent in AR-PIP sets, compared with 11 percent in the proteome. (B) LXXLL-positive prevalence across matched protein-length bins (under 100, 100 to 300, 300 to 600, 600 to 1,200, and over 1,200 amino acids), stratified by compartment. Within every length bin, the cytosolic, microsomal, and nuclear AR-PIPs each carry a lower LXXLL-positive fraction than the whole proteome, confirming the length-controlled depletion observed in Figure 1F is not a modeling artifact. (C) Full detected proteome LXXLL prevalence per compartment (cytosolic n = 5,902, 40.5 percent; microsomal n = 5,509, 42.1 percent; nuclear n = 6,117, 40.6 percent) versus whole UniProt 2023_10 proteome (n = 20,211, 44.7 percent dashed reference). Chi-square p, cyto 1.2 × 10◻⁸, micro 7.8 × 10◻⁴, nuclear 2.7 × 10◻⁸. All three compartmental-detected proteomes are modestly depleted in LXXLL relative to the whole proteome. (D) Length-adjusted LXXLL odds ratio for the nuclear AR-proximal cohort versus the detected nuclear proteome, per androgen time point. The odds ratio is at or below one throughout, at the unliganded baseline it is 0.92 and not significant (open marker), so the early raw prevalence excess reflects protein length rather than LXXLL enrichment, and it becomes progressively depleted with androgen to 0.59 by six hours. (E) Length-controlled logistic regression OR per compartment against compartment-specific detected proteome baseline (this analysis) versus against the whole proteome (from Figure 1F). Cytosolic 0.816 (p = 4.8 × 10◻⁴) versus 0.565 (p = 4.4 × 10◻⁴⁰). Microsomal 0.916 (p = 0.16) versus 0.645 (p = 2.5 × 10◻²⁷). Nuclear 0.683 (p = 1.9 × 10◻¹⁰) versus 0.519 (p = 4.3 × 10◻⁵⁵). Source data, Source Data (sheets Fig_2A, Fig_2B, Fig_2C, Fig_2D, and Fig_2E).

To formally test the direction of depletion after length adjustment, we applied logistic regression of LXXLL+ status on log-transformed protein length and compartment membership. Against the whole-proteome baseline (UniProt 2023_10), the membership odds ratios for LXXLL+ status are 0.565 in cytosolic (p = 4.6 × 10⁻⁴⁰), 0.645 in microsomal (p = 2.5 × 10⁻²⁷), and 0.519 in nuclear (p = 4.5 × 10⁻⁵⁵; Figure 1F). Because the compartmental-detected proteomes are themselves modestly LXXLL-depleted relative to the whole proteome (40.5%, 42.1%, and 40.6% for the cytosolic, microsomal, and nuclear detected proteomes, respectively; Figure 2C), we ran a stricter test using each compartment’s own detected proteome as the reference set. The cytosolic and nuclear depletions remain significant (OR 0.816, p = 4.8 × 10⁻⁴ and OR 0.683, p = 1.9 × 10⁻¹⁰; Figure 2E). The microsomal depletion attenuates to non-significance (OR 0.916, p = 0.16), indicating that the microsomal signal in Figure 1F was largely carried by compartmental composition rather than AR-directed depletion. The nuclear depletion is the strongest single-compartment signal and survives both length and compartment-baseline controls.

Systematic depletion of LXXLL motifs across the AR-PIP universe is biochemically incompatible with high-affinity docking to a single AR LBD surface, which would otherwise be expected to predict enrichment. It is consistent with low-affinity transient engagement at a competitive single surface. We interpret this result through the two-mode framework. Bona fide AR coregulators bind through high-affinity structured-domain contacts at the AR N-terminal domain [20], a mode that does not require LXXLL motifs. The AR AF-2 charge clamp is biochemically receptive to LXXLL motifs in vitro [15,16,17], but AR-specific FQNLF dominates AF-2 occupancy intramolecularly once the N/C interaction commits AR to chromatin [19]. The AF-2 surface therefore transiently and competitively samples the proteome-wide LXXLL pool (Mode 2), and the systematic, length-controlled depletion of LXXLL across the AR-PIP universe is the empirical signature of this competitive mode.

### LXXLL-bearing AR-PIPs in metastatic prostate cancer

We asked whether the 1,981 LXXLL-positive AR-PIPs (union across cytosolic, microsomal, and nuclear compartments; Methods) intersect with prostate cancer (PCa) driver biology. Intersecting with a curated 46-gene PCa driver panel from cBioPortal (TCGA PanCancer Atlas 2018, MSKCC/DFCI 2018, SU2C/PCF 2019; 1,951 pooled samples; Methods) yielded 26 LXXLL-positive AR-PIP drivers, whose alteration frequencies were skewed toward metastatic castration-resistant PCa (mCRPC) relative to localized disease (Figure 3A). AR itself topped the list, dominated by gene amplification (22.3% pooled, 58.8% in mCRPC), the established engine of castration resistance.

**Figure 3.**
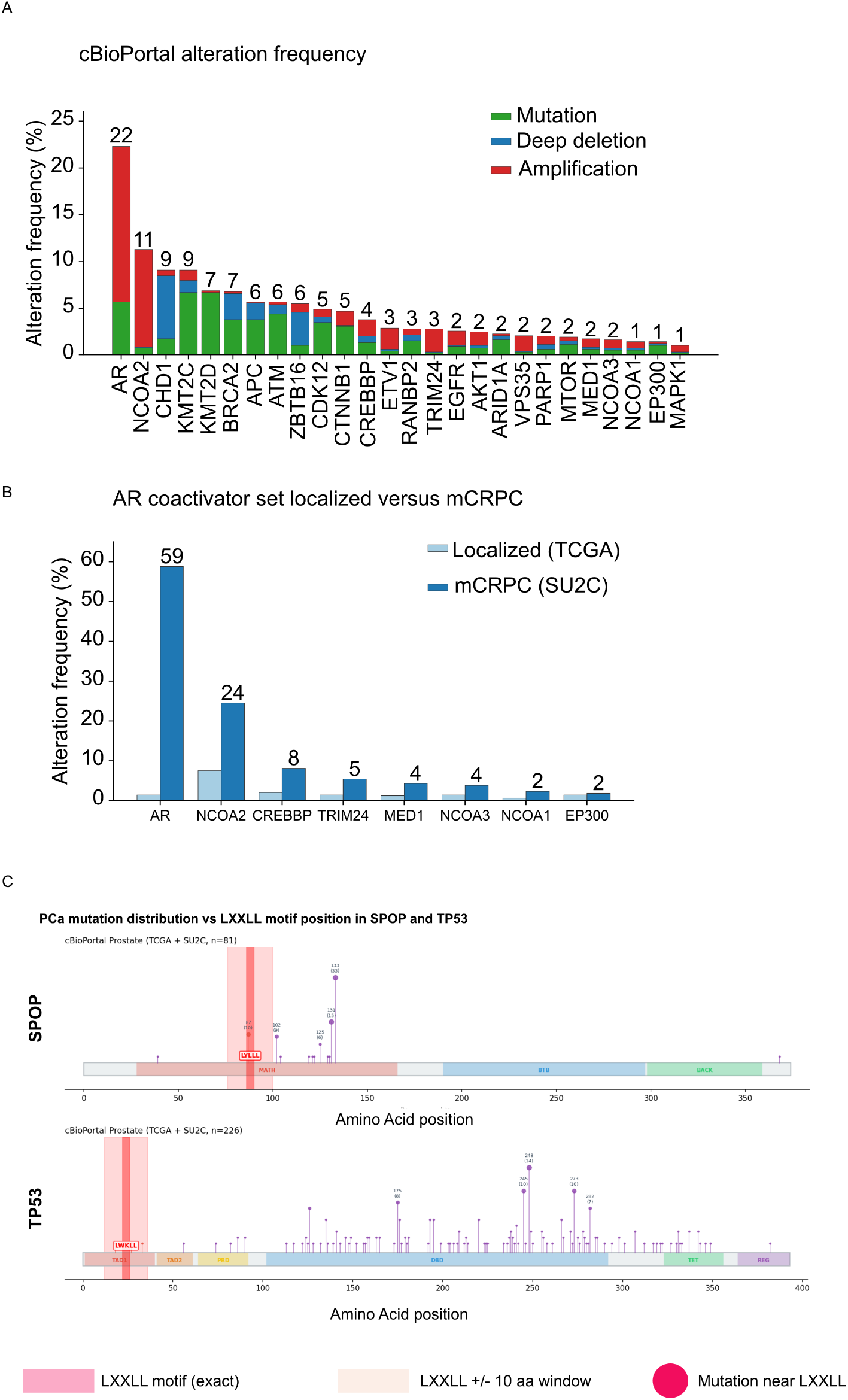
Prostate cancer driver enrichment and SPOP and TP53 mutation clustering relative to LXXLL motifs. (A) Alteration frequency of the 26 LXXLL-positive AR-PIP drivers across three cBioPortal prostate cancer studies (TCGA PanCancer Atlas 2018, MSKCC/DFCI Metastatic 2018 prad_p1000, SU2C/PCF Dream Team 2019; pooled n = 1,951 tumors). (B) Alteration frequency of the canonical AR coactivator cohort (AR, NCOA1, NCOA2, NCOA3, EP300, CREBBP, MED1, TRIM24) skewed toward metastatic castration-resistant prostate cancer (mCRPC). (C) SPOP and TP53 nonsynonymous mutation lollipops from the COSMIC MutantCensus v103 prostate cohort. SPOP mutations cluster around the LYLLL motif at residue 86 within the MATH substrate-recognition domain (top; 1.72-fold enrichment within a 10 amino acid window, p = 4.7 × 10◻⁵). TP53 mutations cluster in the DNA-binding domain at residues 175, 245, 248, 273, 282, distant from the LWKLL motif at TAD1 residues 22 to 26 (bottom). Source data, Source Data (sheets Fig_3A, Fig_3B, and Fig_3C).

Within the 26-gene driver cohort, a structurally diverse but functionally coherent subset of established AR coactivators emerged, comprising AR itself (LTKLL at residue 860 in the ligand-binding domain), NCOA1, NCOA2, NCOA3, p300, CREBBP, MED1, and TRIM24 (Figure 3B). Each carries at least one LXXLL motif, and each is preferentially altered in mCRPC. The convergence of an LXXLL-bearing coactivator cohort onto the mCRPC alteration axis links Mode 2 LXXLL biology to clinical disease progression, going beyond the individual-gene story that AR amplification alone would predict.

At the per-tumor level, LXXLL itself is directly targeted in select drivers. We scanned a 68-gene COSMIC MutantCensus v103 prostate target panel for genes carrying at least one canonical LXXLL motif and at least 20 positioned nonsynonymous mutations, retaining 66, of which 48 are also LXXLL-positive AR-PIPs (Methods). SPOP was the clearest positive. SPOP engages AR directly as a substrate adaptor and targets it for ubiquitination [26,27,28], although it was not recovered in our proximity dataset. PCa SPOP mutations cluster tightly around the LYLLL motif at residue 86 (SPOP UniProt O43791) within the MATH substrate-recognition domain (Figure 3C, top; 1.72-fold enrichment, p = 4.7 × 10⁻⁵), spatially explaining the established mechanism whereby these mutations abolish substrate ubiquitination to stabilize AR. TP53 was likewise analysed from the COSMIC panel and was not recovered in our proximity dataset. Its PCa mutations cluster in the DNA-binding domain at residues 175, 245, 248, 273, and 282, distant from the LWKLL motif at TAD1 residues 22 to 26 (Figure 3C, bottom; 0.25-fold, p = 1), so LXXLL disruption is not the driver mechanism for TP53. Together, the mCRPC-skewed LXXLL-positive coactivator cohort and the SPOP motif-adjacent mutation cluster place LXXLL-bearing AR-PIPs and prostate cancer drivers at the intersection of PCa driver biology and the two-mode framework.

### Functional architectures of LXXLL-positive versus LXXLL-negative AR-PIPs

To dissect what distinguishes LXXLL-positive AR-PIPs functionally from their LXXLL-negative counterparts, we performed g: Profiler enrichment analysis on the six per-compartment protein sets (cytosolic LXXLL+ 1,214 proteins and LXXLL-1,780 proteins; microsomal LXXLL+ 1,518 and LXXLL-1,957; nuclear LXXLL+ 1,424 and LXXLL-1,954) against compartment-specific detected-protein backgrounds (cyto 5,902, microsomal 5,509, nuclear 6,117; Methods). Across all compartments, LXXLL-positive protein sets returned substantially more enriched GO and pathway terms than LXXLL-negative protein sets (cytosolic, 507 versus 145; microsomal, 257 versus 162; nuclear, 724 versus 208 unique terms at BH-corrected p ≤ 0.05; Figure 4A). The asymmetry indicates that LXXLL-positive AR-PIPs occupy functionally coherent biological niches rather than scattering across nonspecific cellular processes.

**Figure 4.**
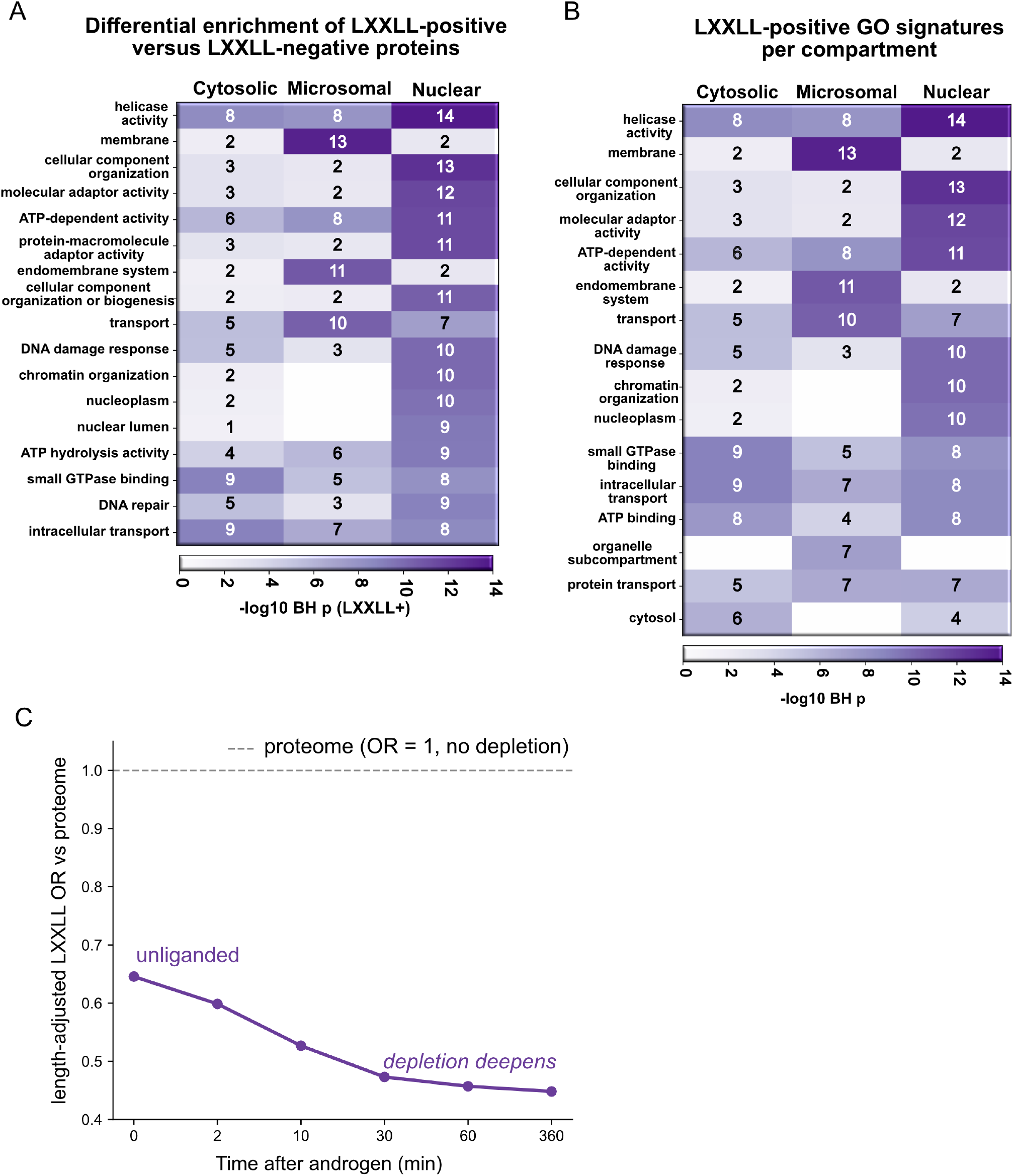
LXXLL-positive AR-PIPs occupy functionally coherent territory, and nuclear LXXLL depletion deepens during chromatin commitment. (A) Differential GO enrichment heatmap for LXXLL-positive versus LXXLL-negative AR-PIPs, showing shared functional core across cytosolic, microsomal, and nuclear compartments. Cells report -log10 Benjamini-Hochberg-adjusted p-values for enrichment in the LXXLL-positive set relative to the LXXLL-negative set. (B) Compartment-distinguishing top GO signatures for LXXLL-positive AR-PIPs. Same purple sequential scale as (A). Cytosolic LXXLL-positive AR-PIPs enrich for small GTPase binding and intracellular transport, microsomal for membrane and endomembrane system, nuclear for adaptor activity and DNA damage response. (C) Length-adjusted LXXLL odds ratio for the nuclear AR-proximal cohort versus the whole proteome, per androgen time point (0, 2, 10, 30, 60, 360 min). The odds ratio is below one (depleted) at every time point and deepens with androgen, from 0.65 at the unliganded baseline to 0.45 at six hours (dashed reference, odds ratio 1). Underlying raw prevalence 47.5, 45.6, 43.3, 39.7, 37.9, 37.1 percent. See also Figure 2. Source data, Source Data (sheets Fig_4A, Fig_4B, and Fig_4C).

Unexpectedly, the LXXLL-positive enrichment signal was not dominated by transcription regulator activity, which would have been the intuitive prediction for a coactivator recognition motif. Instead, the differential signal converged on ATP-dependent activity, helicase function, nucleotide binding, and intracellular transport across all three compartments (Figure 4A). This shared LXXLL-positive functional core is further overlaid with compartment-distinguishing GO terms. Cytosolic LXXLL-positive AR-PIPs feature small GTPase binding and intracellular transport as compartment-distinguishing top hits. Microsomal LXXLL-positive AR-PIPs are associated with the membrane and endomembrane systems. Nuclear LXXLL-positive AR-PIPs feature adaptor activity and DNA damage response (Figure 4B). Each compartment’s LXXLL-positive cohort therefore layers a compartment-appropriate functional axis onto the shared functional core, rather than recapitulating a generic transcription coactivator signature.

The temporal nuclear LXXLL trajectory anchors Mode 2 empirically (Figure 4C). The length-adjusted LXXLL odds ratio for the nuclear AR-proximal cohort relative to the proteome stays below one at every time point and deepens monotonically with androgen, from 0.65 at the unliganded baseline (0 minutes) to 0.45 at six hours, with the underlying raw prevalence falling in parallel from 47.5% to 37.1% (45.6%, 43.3%, 39.7%, and 37.9% at 2, 10, 30, and 60 minutes). Controlled against the flat nuclear-detected proteome baseline, the cohort is not LXXLL-enriched even at the unliganded baseline (odds ratio 0.92, not significant) and becomes progressively depleted with androgen (odds ratio 0.59 by six hours; Figure 2D), so the decline is AR-directed rather than compositional drift and directly reflects the Mode 2-to-Mode 1 transition. By 360 minutes, AR is chromatin-committed via the FQNLF N/C interaction, displacing the LXXLL-positive cohort. Mode 2 transient LXXLL engagement at the AR AF-2 charge clamp is permissive during cytoplasmic-to-nuclear transit. Still, it is outcompeted by intramolecular FQNLF dominance once the receptor commits to its DNA-bound state.

### AR-V7 as a natural AF-2-deletion test of the two-mode framework

The two-mode framework predicts that loss of the AF-2 charge clamp should selectively strip Mode 2 LXXLL-positive partners while leaving Mode 1 structured-domain partners intact. The AR-V7 splice variant, which terminates before the ligand-binding domain and therefore lacks the AF-2 surface entirely [4], provides a natural human experiment to test this prediction. If Mode 2 LXXLL engagement is AF-2 dependent, the proximal proteome of AR-V7 should be depleted of LXXLL-positive partners relative to full-length AR, and the partners lost from AR-V7 should be LXXLL-enriched.

To test this directly, we compared the published AR-V7 proximal proteome from Adamson et al (2023; FLAG-APEX2-AR-V7 in CWR22Rv1 castration-resistant cells; 463 proteins after HGNC normalization) with our full-length AR-PIP universe (4,751 proteins). Three protein sets emerged. The shared set (Adamson AR-V7 intersecting with our full-length AR-PIPs) contained 343 proteins, 37.9% of which were LXXLL-positive. The full-length-only set (our AR-PIPs not in Adamson AR-V7) contained 4,408 proteins, 42.0% of which were LXXLL-positive. The AR-V7-only set (Adamson hits not in our AR-PIPs) contained 120 proteins, 26.5% of which were LXXLL-positive (Figure 5A). The AR-V7 interactome itself was 35.0% LXXLL-positive, depleted relative to the human proteome baseline of 44.7% and to our full-length AR-PIP union at 41.7%. These results are consistent with the loss of an LXXLL-receptive surface in AR-V7.

**Figure 5.**
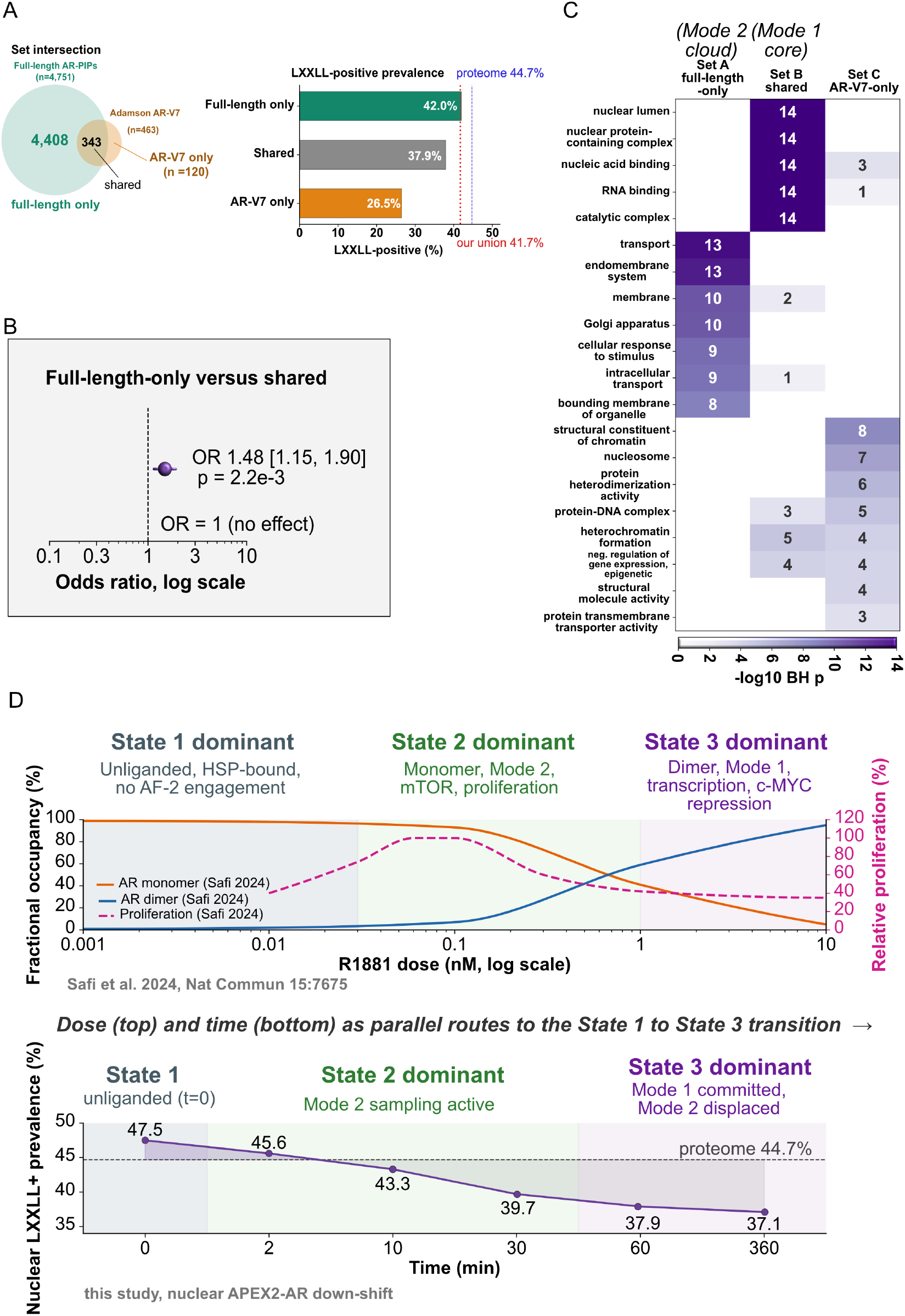
AR-V7 lacks AF-2 and loses LXXLL-enriched Mode 2 partners. (A) Set intersection of the Adamson et al 2023 AR-V7 proximal proteome (n = 463 proteins) with our full-length AR-PIP universe (n = 4,751 proteins). Full-length only n = 4,408, shared n = 343, AR-V7 only n = 120. Right panel, LXXLL-positive prevalence per set (42.0, 37.9, 26.5 percent). Whole-proteome 44.7 percent dashed reference, our AR-PIP union 41.7 percent dotted reference. (B) Length-controlled logistic regression odds ratio comparing full-length-only against shared set (reference), adjusted for log protein length. OR 1.48, 95 percent CI 1.15 to 1.90, p = 2.2 × 10◻³. Vertical dashed line at OR = 1 marks no effect. (C) g:Profiler GO enrichment heatmap for LXXLL-positive proteins per set. Set A full-length-only (Mode 2 cloud, n = 1,851). Set B shared (Mode 1 core, n = 130). Set C AR-V7 only (n = 31). Cells report -log10 Benjamini-Hochberg adjusted p. Union of detected proteomes used as background. (D) Three-state framework overlaid on parallel dose [29] and time (this study, lower) axes. Upper, AR monomer occupancy (orange), AR dimer occupancy (blue), and relative proliferation (dashed magenta) versus R1881 dose. Lower, nuclear LXXLL-positive prevalence per time point (this study, purple), proteome 44.7 percent dashed reference. Three shaded regions on each axis mark State 1 (blue-gray, unliganded, HSP-bound), State 2 (green, monomer, Mode 2 sampling), State 3 (purple, dimer on chromatin, Mode 1 committed). See also Figure 2. Source data, Source Data (sheets Fig_5A through Fig_5D); the dose axis in (D) is replotted from Safi et al 2024.

If protein length is accounted for, the signal strengthens. This is because LXXLL is a five-residue motif that can arise anywhere along a sequence, and longer proteins contain LXXLL more often by chance. Any group biased toward larger members appears artificially LXXLL-positive in raw prevalence. This confound is present here. The shared set (median 621 amino acids) is longer than the full-length-only set (median 575 amino acids), inflating the shared set’s raw LXXLL rate and narrowing the apparent gap. To remove this bias, we modeled per-protein LXXLL positivity by logistic regression with log protein length as a covariate. At matched length, a protein in the full-length-only set is 1.48-fold more likely to contain an LXXLL motif than a protein in the shared set (95% CI 1.15 to 1.90, p = 2.2 × 10⁻³; Figure 5B). The raw 42.0% versus 37.9% comparison, therefore, understates the loss of LXXLL enrichment upon AF-2 deletion. AF-2 loss preferentially discards LXXLL-positive proximal partners, the empirical signature Mode 2 predicts. The shared set represents the LXXLL-depleted Mode 1 core that bona fide coregulators access at the AR N-terminal domain, and the full-length-only set represents the LXXLL-enriched Mode 2 cloud that requires AF-2. Both studies rely on APEX2 proximity biotinylation, matching the labeling chemistry and radius across datasets. Beyond that shared framework, cell-line background and bait construct diverge. Adamson used FLAG-APEX2-AR-V7 in castration-resistant CWR22Rv1 cells that express full-length AR and AR variants. We used APEX2-AR in androgen-responsive LNCaP cells that express only full-length AR. The shared and full-length-only sets therefore reflect these background differences, as well as the intended AR-V7-versus-full-length contrast. The length-controlled odds ratio of 1.48 supports the two-mode prediction under these constraints. A matched comparison of full-length AR and AR-V7 in CWR22Rv1 cells would provide a definitive test.

We next asked what biology distinguishes the three sets. g: Profiler enrichment on each LXXLL-positive subset (union of detected proteomes as background; Methods) revealed nearly non-overlapping signatures (Figure 5C). Set A (Mode 2 cloud, n = 1,851) enriched for transport, endomembrane system, membrane, Golgi apparatus, and intracellular transport (BH-p ≤ 10⁻⁸), the trafficking biology sampled by AF-2 during cytosolic-to-nuclear transit. Set B (Mode 1 core, n = 130) enriched for nuclear lumen, nuclear protein-containing complex, nucleic acid binding, RNA binding, and catalytic complex (all BH-p ≤ 10⁻¹⁴), retaining the structured coregulator biology without AF-2. Set C (AR-V7-only, n = 31) was enriched for structural constituents of chromatin, nucleosomes, protein heterodimerization activity, and heterochromatin formation (BH-p 10⁻³ to 10⁻⁸), a chromatin-remodeling signature specific to Adamson’s CWR22Rv1 AR-V7 dataset. The near block-diagonal functional structure reinforces what the length-controlled OR 1.48 shows statistically. AF-2 loss preferentially discards the LXXLL Mode 2 cloud, retains the Mode 1 core, and reveals chromatin-remodeler biology outside the full-length AR sampling window.

Together, the AR-V7 evidence integrates with the temporal nuclear LXXLL trajectory (Figure 4C) and prior dose-response modeling of AR monomer and dimer partitioning into a coherent three-state framework [29]. Dose and time represent parallel empirical routes through the same State 1 (unliganded, HSP-bound), State 2 (liganded monomer, Mode 2 sampling active), and State 3 (liganded dimer on chromatin, Mode 1 committed, Mode 2 displaced) transition (Figure 5D).

### Conservation of the LXXLL coactivator core across the nuclear-receptor family

The two-mode framework predicts that the LXXLL coactivator vocabulary should be conserved across nuclear receptors that share the AF-2 charge-clamp architecture. To test this at the interactome level beyond structural conservation, we intersected the published high-confidence GR interactome from Lempiäinen et al (2017; dexamethasone-agonist arm, HEK293 cells, BioID, SAINT BFDR ≤ 0.01, NR3C1 bait excluded; 24 partners) with our LNCaP AR-PIP universe. 20 of 24 (83.3%) Lempiäinen GR partners are recovered in our AR proximal proteome, and 16 of these 20 (80%) are LXXLL-positive in both datasets. The cross-lab, cross-method, cross-cell-line shared set comprises canonical nuclear-receptor coactivators (NCOA2, NCOA3, NCOA6, NCOR1, MED1) and chromatin-remodeling complexes (SWI/SNF components ARID1A, ARID1B, SMARCA4, SMARCC1, SMARCC2; KDM1A; KMT2D) (Figures 6A-6C). The concordance extends Lempiäinen et al’s within-experiment AR/GR interactome overlap to an independent laboratory, cell system, and proximity-labeling chemistry, establishing that the LXXLL-engaging coactivator core is a shared family engine rather than receptor-specific machinery.

**Figure 6.**
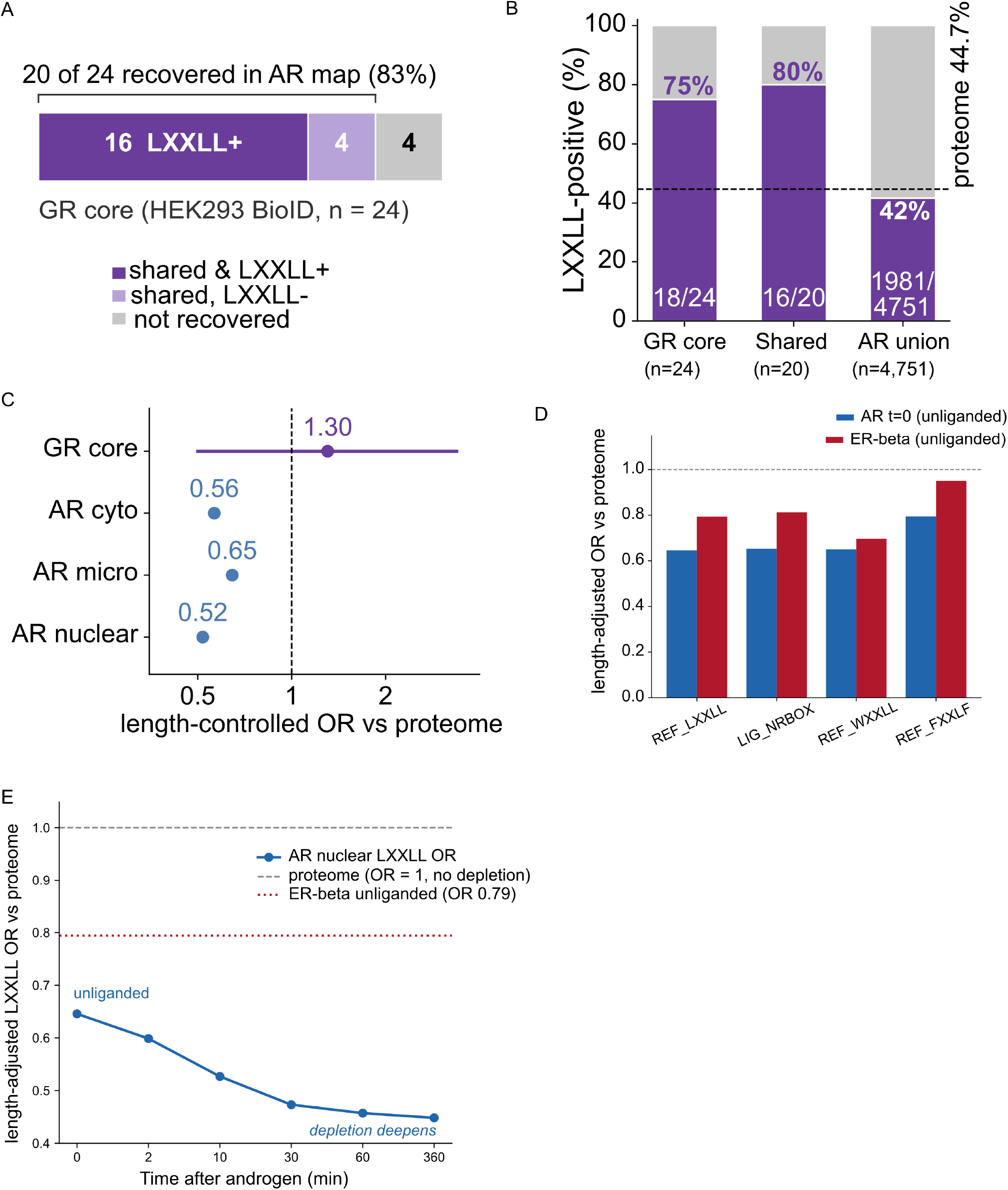
The LXXLL coactivator core is a shared nuclear-receptor family engine. (A) GR-core containment. Of 24 high-confidence GR proximal partners (dexamethasone arm, HEK293 cells, BioID, SAINT BFDR ≤ 0.01, NR3C1 bait excluded)[5], 20 (83%) are recovered in the LNCaP AR-PIP universe (n = 4,751), and 16 of the 20 shared partners (80%) are LXXLL-positive in both datasets. The shared core includes canonical nuclear receptor coactivators (NCOA2, NCOA3, NCOA6, NCOR1, MED1) and chromatin-remodeling complexes (ARID1A, ARID1B, SMARCA4, SMARCC1, SMARCC2, KDM1A, KMT2D); the four not recovered are ACACB (a BioID biotinylated carboxylase background protein), ARID3B, CPVL, and TUBB4A. (B) LXXLL-positive prevalence across interactome depth. The stable shared coactivator core is LXXLL-enriched (GR core 75.0%, 18 of 24; AR-GR shared set 80.0%, 16 of 20) relative to the whole-proteome baseline (44.7%), whereas the broad AR-PIP union sampled by APEX2 is LXXLL-depleted (41.7%, 1,981 of 4,751). (C) Length-controlled LXXLL membership odds ratios versus the proteome. The GR-core odds ratio (1.30, 95% CI 0.50 to 3.38) is not significant, whereas all three AR compartments are significantly depleted (cytosolic 0.565, microsomal 0.645, nuclear 0.519); LXXLL depletion is therefore a property of interactome depth, not receptor identity. (D) Cross-receptor coactivator-motif depletion. Length-adjusted odds ratios versus the proteome for coactivator-recognition motif classes in unliganded AR (t = 0) and the independent unliganded nuclear ER-β interactome [30]. Both receptors are depleted for LXXLL, the NR-box, and WXXLL relative to the proteome. (E) Length-adjusted LXXLL depletion deepens across the androgen time course. The length-adjusted LXXLL odds ratio for the nuclear AR proximal interactome, versus the proteome, remains below one at every time point, declining from 0.65 (unliganded) to 0.45 (six hours after androgen); the unliganded ER-β interactome (odds ratio 0.79, dotted line) sits at the same depleted, pre-androgen level, above the AR trajectory but below the no-depletion reference (odds ratio 1, dashed line). Source data, Source Data (sheet Fig_6); ER-β odds ratio derived from Giurato et al 2018 (ProteomeXchange PXD006720).

An independent test in a second nuclear receptor reinforced this conclusion. We compared the AR-PIP length-controlled motif analysis with the nuclear ER-β interactome reported by Giurato et al.[30] (an RNA-mediated ER-β interactome purified from MCF-7 breast cancer cell nuclei by label-free quantitative proteomics). In the matched unliganded state, the ER-β interactome showed the same length-controlled depletion of coactivator-recognition motifs as unliganded AR. The length-adjusted LXXLL odds ratio relative to the proteome was 0.79 for ER-β and 0.65 for AR at t = 0, with concordant depletion of the NR-box and WXXLL motif classes (Figure 6D). Consistent with the AR temporal trajectory, the length-adjusted LXXLL odds ratio remained below one at every time point and deepened with androgen exposure, from 0.65 in the unliganded state to 0.45 by six hours; the unliganded ER-β interactome (odds ratio 0.79) sat at the same depleted, pre-androgen level (Figure 6E). The convergence of an independent receptor, laboratory, cell system, and purification chemistry indicates that length-controlled coactivator-motif depletion is a shared property of the nuclear-receptor proximal interactome rather than a feature specific to AR.

## Discussion

By integrating the cytosolic, microsomal, and nuclear proximal proteomes of LNCaP prostate cancer cells, we resolve 4,751 AR-proximal interacting proteins, four to five times the curated AR-interactome. The breadth of this proximal universe poses a structural question. What features of the AR ligand-binding domain permit a single receptor to sample so many partners at stoichiometrically meaningful levels? Length-controlled depletion of LXXLL across all compartments provides an answer. The pattern is consistent with low-affinity, transient engagement rather than high-affinity docking, and it predicts a per-time-point nuclear LXXLL trajectory that declines from 47.5% at baseline to 37.1% at six hours as AR commits to chromatin.

Three lines of evidence support the two-mode framework (Figures 7A and 7B). Full-length cryo-EM [19,20] shows that bona fide AR coregulators bind the AR N-terminal domain through structured bHLH/PAS, S/T, and HAT contacts independent of LXXLL, so Mode 1 cannot drive LXXLL enrichment. The AR AF-2 charge clamp is biochemically receptive to LXXLL motifs [15,16,17] but is sequestered by intramolecular FQNLF once AR commits to chromatin [19,31], thereby making Mode 2 sampling commitment-gated. The AR-V7 splice variant, which lacks AF-2 entirely, preferentially retains LXXLL-depleted Mode 1 partners and loses the LXXLL-enriched Mode 2 cohort at length-controlled OR 1.48. Together, these identify Mode 2 as a transient, competitive, charge-clamp-gated sampling of the proteome-wide LXXLL pool, and length-controlled depletion as its empirical signature.

**Figure 7.**
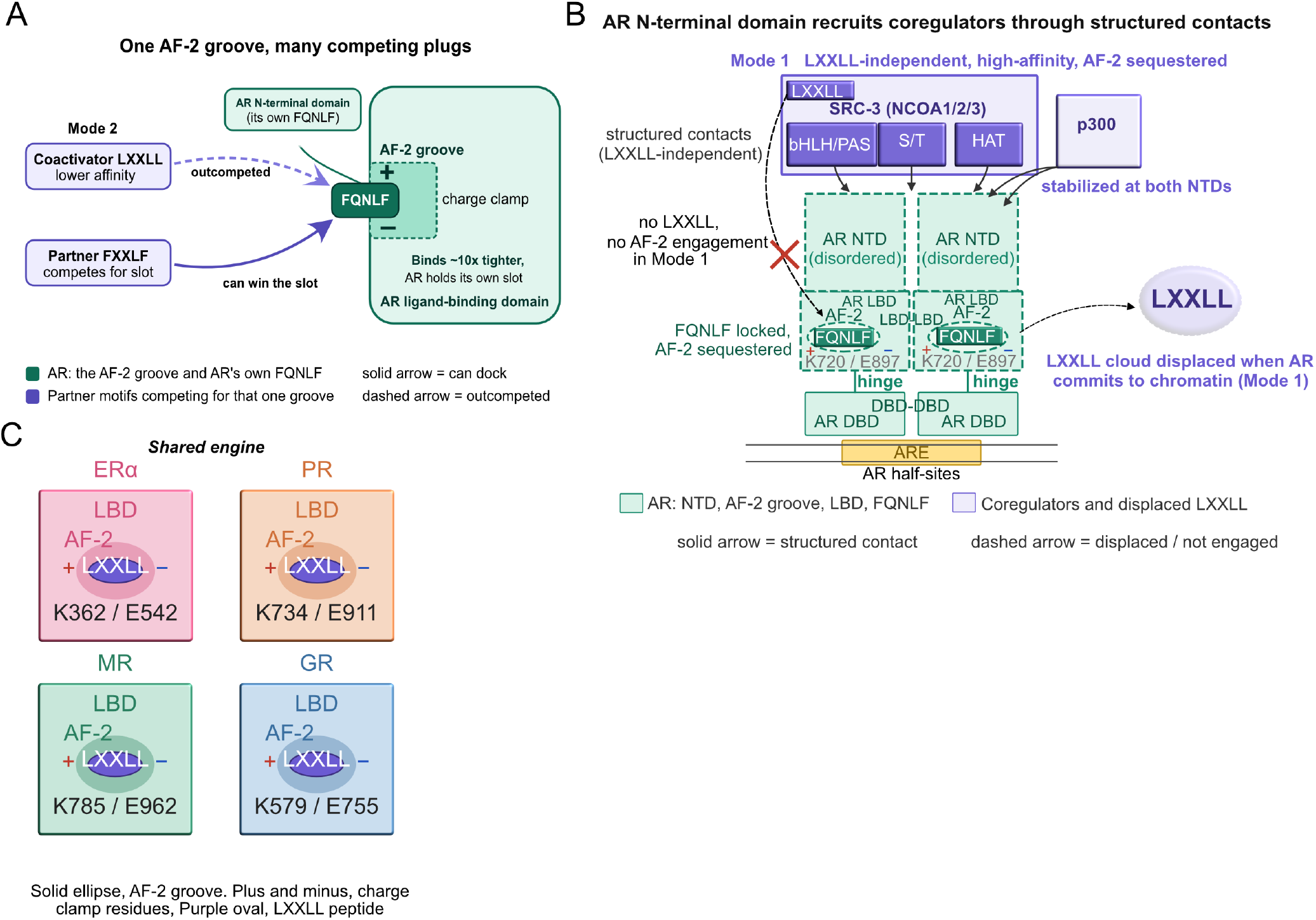
Two-mode framework and nuclear-receptor family generalization. (A) Mode 2, transient LXXLL sampling at the AR AF-2 charge clamp. AR’s intrinsic FQNLF motif dominates AF-2 occupancy (approximately 10-fold higher affinity than coactivator LXXLL). Partner FXXLF motifs can compete for the slot. Coactivator LXXLL is outcompeted. Green, AR components. Purple, partner motifs. Solid arrow, docks. Dashed arrow, outcompeted. (B) Mode 1, structured-domain coregulator recruitment at the AR N-terminal domain in the chromatin-committed AR dimer. SRC-3 (NCOA1/2/3) engages the AR NTD through bHLH/PAS, S/T, and HAT domain contacts, independently of LXXLL. p300 stabilized at both NTDs. AR dimer bound to ARE, DBD-DBD interface at the ARE, LBD-LBD interface between LBDs, intramolecular FQNLF locked in each AF-2 groove sequestering the charge clamp (K720, E897). LXXLL cloud displaced. Solid arrow, structured contact. Dashed arrow displaced or not engaged. Cryo-EM geometry follows Yu et al 2020. (C) The nuclear receptor family AF-2 charge-clamp architecture is conserved across the estrogen receptor alpha (ERα), progesterone receptor (PR), mineralocorticoid receptor (MR), and glucocorticoid receptor (GR). Each LBD schematic shows the AF-2 groove, charge clamp (plus and minus), and an in-groove LXXLL peptide (purple). Residue callouts, ERα K362/E542, PR K734/E911, MR K785/E962, GR K579/E755 (UniProt canonical). AR K720/E897 in Figure 7B follows the standard literature 22-glutamine reference polyQ numbering; see Methods. Figure 7 is a schematic summary; no source data are associated with this figure.

The AR proximity data are consistent with a shared-engine model of nuclear-receptor interactome organization (Figure 7C). Because the LXXLL coactivator core is shared between AR and GR [5], LXXLL cannot itself determine receptor specificity. It is a common engine. Specificity, the property that assigns each receptor to its own transcriptional context and prevents AR and GR from colliding over the shared coactivator pool in cells that respond to both hormones, is instead encoded in the receptor-specific and largely non-LXXLL transient interactions of the Mode 2 cloud. This framing explains why LXXLL motifs are depleted throughout the broad proximal AR proteome. The bulk of each receptor’s context-defining partners engage through receptor-specific SLiMs rather than the shared LXXLL vocabulary. In prostate cancer, AR and GR avoid substrate competition through reciprocal deployment. AR normally suppresses GR expression, and GR emerges under AR blockade to activate a similar but distinguishable target gene program [32]. Similar reflects the shared LXXLL coactivator core; distinguishable reflects receptor-specific Mode 2 routing. Estrogen, progesterone, glucocorticoid, and mineralocorticoid receptors share the AF-2 charge-clamp architecture and the LXXLL recognition vocabulary [13,33], and analogous Mode 1 structured-domain surfaces have been reported for their N-terminal domains. Together with the companion cytosolic-microsomal [9] and nuclear [10] AR-PIN atlases, the shared-engine framework establishes a conceptual scaffold for how a common family recognition vocabulary accommodates diverse receptor-context specificity across the nuclear receptor superfamily.

### Limitations of this study

Several limitations merit consideration. First, the AR-V7 comparison relies on Adamson et al.[4], who used FLAG-APEX2-AR-V7 in castration-resistant CWR22Rv1 cells, whereas our AR-PIP universe uses APEX2-AR in androgen-responsive LNCaP cells. Labeling chemistry is shared, but differences in cell-line background and bait construct contribute to the boundaries between the shared and full-length-only sets, alongside the AR-V7 versus full-length contrast. The length-controlled odds ratio of 1.48 (p = 2.2 × 10⁻³) supports the two-mode prediction under these constraints. A matched cell-line comparison of full-length AR and AR-V7 would provide a definitive test. Second, the integrated dataset is a discovery resource. Beyond the AR-V7 natural ΔAF-2 experiment, this study does not directly perturb the AR AF-2 charge clamp itself; structure-guided point mutations at the charge-clamp residues (Lys720, Glu897) will be needed to definitively test the two-mode framework. Third, our proximity-labeling data cannot distinguish independent Mode 1 and Mode 2 coregulator populations from a tethered-handoff model in which the same coregulator engages both surfaces sequentially, first through LXXLL contacts at AF-2 in a prebound state and then through structured N-terminal domain contacts once intramolecular FQNLF locks the AF-2 groove upon agonist binding. Direct biochemical and structural work will be needed to distinguish these scenarios.

## Methods

Unless otherwise stated, all analyses used HGNC-canonical gene aggregation and a fixed random seed, set.seed(20260601). All results were generated from current pipeline outputs.

### Proteomic Data Analysis

This study uses the AR-PIP proximal-interaction datasets derived from the differential-abundance pipeline detailed in the companion cytosolic-microsomal [9] and nuclear [10] manuscripts. In brief, DIA-NN [34] protein-group quantifications were processed through variance-stabilizing normalization [35], Perseus-style downshift imputation [36] (width 0.3 σ, downshift 1.8 σ, seed set.seed(20260601)), and limma moderated t-tests [37] with Benjamini-Hochberg false discovery rate correction. AR-PIP status was defined by adjusted p ≤ 0.05 and log2 fold-change > 1 relative to the no-H2O2 background control. The R statistical pipeline is available at https://github.com/mewrightlab/DIA-Toolkit (v1.0.0-AR-PINs).

### MotifHunter Sequence Scan

MotifHunter, a sequence-scan algorithm developed for this study (see the Key Resources supplementary table for repository URL), was used to identify occurrences of the LXXLL, FXXLF, and WXXLL motifs in canonical Swiss-Prot UniProt sequences (UniProt 2023_10 reviewed entries, n = 20,211). For each AR-PIP, the canonical protein sequence was scanned for all matches to the regular expression patterns L[A-Z][A-Z]LL, F[A-Z][A-Z]LF, and W[A-Z][A-Z]LL. Each protein was scored as motif-positive if it contained at least one match. Compartmental AR-PIP sets were split by motif content, and chi-square tests against the UniProt proteome baseline were performed for each compartment.

During the preparation of this manuscript, the authors used Claude (Anthropic) to assist in writing C++ code for the Motif Hunter program. All code was validated and the authors take full responsibility for the software.

### Protein-Length Logistic Regression

To control for protein-length confounding in the LXXLL prevalence comparisons, logistic regression was used to model LXXLL-positive status as a function of the natural logarithm of canonical protein length and a categorical compartment factor. Membership odds ratios for the cytosolic, microsomal, and nuclear AR-PIP sets were calculated against the UniProt 2023_10 proteome baseline. Wald p-values, 95% confidence intervals, and per-bin LXXLL prevalence within five length bins (less than 100, 100 to 300, 300 to 600, 600 to 1200, and greater than 1200 amino acids) were generated to support the AR-PIP-wise length distribution analysis.

### cBioPortal Genomic Analysis

Alteration frequency analysis was performed in cBioPortal across three prostate cancer studies (TCGA PanCancer Atlas 2018, MSKCC/DFCI Metastatic 2018 prad_p1000, and SU2C/PCF Dream Team 2019; 1,951 pooled samples). The unified LXXLL-positive AR-PIP set (union across the three compartments, 1,981 proteins) was intersected with a curated 46-gene prostate cancer driver panel (Supplementary Data 1, sheet Driver_Panel), yielding 26 LXXLL-positive AR-PIP drivers. Alteration types included nonsynonymous point mutations and discrete copy number alterations. Per-gene alteration frequencies were stratified by disease state (localized versus metastatic castration-resistant prostate cancer).

### COSMIC Mutation Window Analysis

Somatic mutation positions were retrieved from a 68-gene COSMIC MutantCensus v103 prostate target panel. For each gene in that panel carrying at least one canonical LXXLL motif and at least 20 positioned nonsynonymous mutations, the fold-enrichment of mutations within a 25-residue window spanning each LXXLL motif and the 10 residues flanking it on either side (motif start minus 10 through motif start plus 14) was calculated relative to expectation, given the total protein length and total mutation count. Motif positions were re-derived from canonical UniProt sequences rather than from curated motif annotations. Significance was assessed by one-sided binomial testing. Per-sample motif-spanning mutations were also tallied for selected drivers (TP53, SPOP) to support the per-tumor motif-disruption analysis.

### AR-V7 vs Full-Length AR Comparison

The Adamson et al (2023) Supplementary Table 4 AR-V7 APEX2 proximal proteome (463 HGNC-normalized proteins, org.Hs.eg.db 3.18.0) was intersected with the full-length AR-PIP universe (4,751 proteins; union of cytosolic, microsomal, and nuclear AR-PIPs). Three protein sets were defined and shared (intersection of Adamson AR-V7 and our full-length AR-PIPs, 343 proteins). Full-length only (our AR-PIPs not in Adamson, 4,408 proteins). AR-V7 only (Adamson hits not in our full-length AR-PIPs, 120 proteins). LXXLL-positive fraction was computed per set. Logistic regression of LXXLL-positive status on log-length and set membership was used to compute the length-controlled odds ratio for full-length-only versus shared-set membership.

### Cross-Compartment Set Operations

The full LNCaP AR-PIP universe was assembled as the HGNC-canonical union of the cytosolic, microsomal, and nuclear AR-PIP sets from the companion atlases [9,10]. Pairwise and three-way intersections were computed for the UpSet plot panel. Per-compartment expansion factors were calculated as the ratio of compartmental AR-PIP union size to the detected fraction of the 989-member curated AR-IP catalog in that compartment.

Key resources, including antibodies, cell lines, chemicals, kits, and software, are provided in Supplementary Table 1 (Key Resources).

### Data availability

The mass spectrometry proteomics data underlying this study were generated jointly with the companion cytosolic-microsomal and nuclear manuscripts and have been deposited to the ProteomeXchange Consortium via the PRIDE partner repository [38] with the dataset identifier PXD081449. Source data for all main and supplementary figures are provided in the Source Data file, and the AR-PIP universe, AR-V7 partitions, and driver panel are provided in Supplementary Data 1. All other data reported in this paper will be shared by the corresponding author upon request.

### Code availability

Analysis code, including MotifHunter (the LXXLL, FXXLF, and WXXLL sequence-scanning algorithm), the protein-length-controlled logistic regression pipeline, the AR-V7 set-comparison workflow, and the shared-engine GR interactome overlap analysis is deposited at https://github.com/mewrightlab/DIA-Toolkit, and MotifHunter is additionally available at https://github.com/UWPR/CGI_tools/tree/main/motif_search/. The repository is private during peer review, and reviewer access is available from the corresponding author. Upon manuscript acceptance, the repository will be made publicly accessible, a release tag (v1.0.0-AR-PINs) will be cut, and an archival Zenodo DOI ([Zenodo DOI pending]) will be minted and added to the published record. The pipeline was developed and tested on R 4.3.2 (Bioconductor 3.18).

## Acknowledgements

This work was supported by the National Institutes of Health (R01GM143399).

## Author contributions

J.K.E. developed MotifHunter, the LXXLL, FXXLF, and WXXLL sequence-scanning algorithm, and contributed to the computational analysis of the mass spectrometry data. M.E.W. conceived and designed the study, performed the compartment-resolved motif and enrichment analyses, interpreted the data, and wrote the manuscript with input from all authors. L.R. contributed to data interpretation and edited the manuscript. M.E.W. supervised the project and secured funding. All authors reviewed and approved the final manuscript.

## Use of generative AI

During the preparation of this work, the authors used Claude (Anthropic) to assist with manuscript drafting, editing, formatting, and reference management, and to assist in writing the C++ code for the MotifHunter program. All code was validated and all computational outputs were independently reviewed by the authors against the primary data. The authors reviewed and edited all AI-assisted content as needed and take full responsibility for the content of the publication.

